# Peroxisomal tail-anchored proteins do not reach peroxisomes via ER, instead mitochondria can be involved

**DOI:** 10.1101/2023.05.22.541719

**Authors:** Tamara Somborac, Güleycan Lutfullahoglu Bal, Kaneez Fatima, Helena Vihinen, Anja Paatero, Eija Jokitalo, Ville O Paavilainen, Svetlana Konovalova

## Abstract

Peroxisomes are membrane-enclosed organelles with important roles in fatty acid breakdown, glycolysis, and biosynthesis of sterols and ether lipids. Defects in peroxisome biogenesis result in severe neurological diseases, such as Zellweger syndrome, neonatal adrenoleukodystrophy, infantile Refsum disease, and myelopathies. However, many aspects of peroxisomal biogenesis are not well understood. Here we investigated delivery of tail-anchored (TA) proteins to peroxisomes in mammalian cells. Using glycosylation assays we showed that peroxisomal TA proteins do not enter ER in both WT and peroxisome-lacking cells. We observed that in cells lacking the essential peroxisome biogenesis factor, PEX19, peroxisomal TA proteins localize mainly to mitochondria. However, in PEX3 KO cells, which lack peroxisomes as well, the endogenous TA protein, ACBD5, does not target mitochondria, suggesting that PEX3 plays an important role in targeting of peroxisomal TA proteins to mitochondria. Finally, to investigate peroxisomal TA protein targeting in cells with fully functional peroxisomes we used a proximity biotinylation approach. We showed that while ER-targeted TA construct was exclusively inserted into the endoplasmic reticulum (ER), peroxisome-targeted TA construct was inserted to both peroxisomes and mitochondria. Thus, in contrast to previous studies, our data suggest that peroxisomal TA proteins do not insert to the ER prior to their delivery to peroxisomes. Instead, mitochondria can play a role in the targeting of TA proteins to peroxisomes.

## INTRODUCTION

Peroxisomes are single membrane-bounded organelles with essential metabolic functions in human cells. Peroxisomes are crucial in lipid metabolism, catabolism of very long chain fatty acids, biosynthesis of sterols, and ether lipids. Defects in peroxisome biogenesis result in severe neurological diseases, such as Zellweger syndrome, neonatal adrenoleukodystrophy, infantile Refsum disease, and myelopathies ^1^. De novo peroxisomal biogenesis includes assembly of peroxisomal membrane proteins and subsequent import of soluble proteins into peroxisomal matrix ^2, 3^ . Although the main components and pathways of peroxisomal matrix protein import have been successfully identified ^4–6^ how peroxisomal membrane proteins (PMPs) are assembled remains unclear ^7, 8^.

A small portion of PMPs are embedded in the peroxisomal membrane by their C-terminal transmembrane segment, termed the tail-anchor (TA) that can direct newly synthesized TA proteins to different subcellular compartments. A combination of hydrophobicity of transmembrane domain and tail charge of TA proteins may play a role in the specific targeting of TA proteins to peroxisomes ^9^. Tail-anchor proteins (TA) carry out important functions in cell metabolism ^10^ and several disorders are linked to defective targeting of these proteins ^11, 12^. Studies on PMP targeting in mammalian cells have suggested different modes of delivery to peroxisomes: a direct delivery from cytosol to pre-existing organelles or/and ER dependent trafficking. In the direct delivery model, PMPs reach peroxisomes without ER involvement by using essential peroxisome biogenesis factors ^13, 14^. On the other hand, the indirect pathway suggests that PMPs are initially inserted into the ER membrane before being sorted to peroxisomes. It has been demonstrated that in the absence of mature peroxisomes many PMPs accumulate at the ER membrane ^15, 16^. Several distinct mechanisms were suggested to play the role in this process, such as involvement of Sec61 translocon ^17^ and essential peroxins in the trafficking of TA proteins through the ER.

Two highly conserved peroxins Pex19 and Pex3, are thought to be the key elements of PMP trafficking in cells ^18^. While Pex19 is mainly a cytosolic chaperone, Pex3 acts as a membrane docking factor for protein insertion ^10^. Deletion of one of these peroxins results in cells devoid of mature peroxisomes. While previous studies focused on the interplay between essential biogenesis factors, no detailed biochemical studies have investigated TA delivery to peroxisomes in mammalian cells.

We studied the pathway of TA targeting to peroxisomes using biochemical assays and microscopy. We took advantage of YgiM(TA), a well-characterized bacteria-derived tail-anchored protein (TA) targeted specifically to peroxisomes in mammalian cells ^19^. Its small size and lack of function-directed interactions within eukaryotic cells make it a versatile tool to investigate protein targeting to peroxisomal membrane. Our data suggest that peroxisomal TA proteins do not insert ER prior to their delivery to peroxisomes. Instead, mitochondria can play a role in the targeting of TA proteins to peroxisomes.

## RESULTS

### Peroxisomal TA proteins do not transit through the ER in mammalian cells

We rationalized that if peroxisomal TA proteins become exposed to the ER lumen at some point during the delivery to mature peroxisomes, then in the absence of peroxisomes, peroxisomal TA proteins could be detectable at the ER. Therefore, we generated HEK293T cell lines lacking peroxisomes by deletion of the essential peroxisomal biogenesis factors, PEX3 or PEX19, using CRISPR Cas9 editing. Immunoblotting of the resulting cell lines showed the specific deletion of either PEX3 or PEX19 (Fig. 1a). As expected from previous research ^13, 14^, confocal microscopy revealed lack of peroxisomes in both cell lines as assessed by immunofluorescence labeling of catalase (Fig. 1b).

**Fig. 1.**
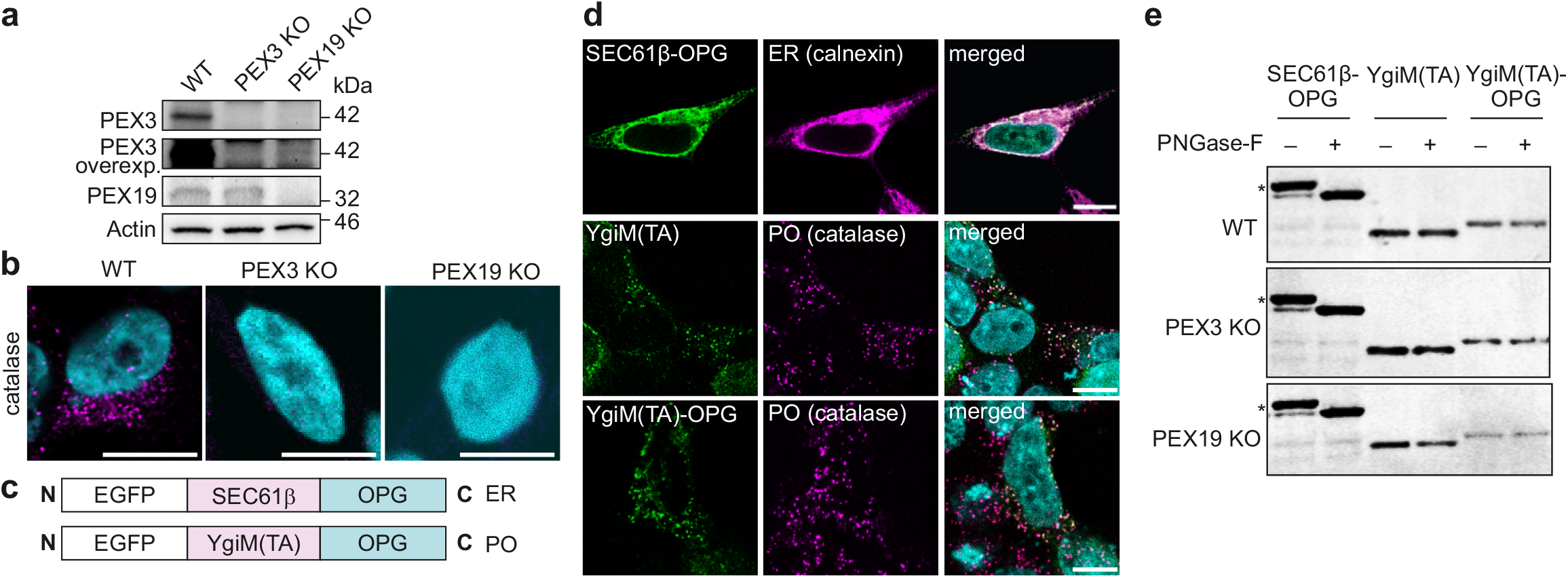
Peroxisomal TA proteins do not transit through the ER in mammalian cells. (a) Immunoblotting analysis of PEX3 KO or PEX19 KO HEK293T cells generated by CRISPR Cas9 approach. Actin was used as a loading control. (b) Confocal immunofluorescence analysis of peroxisomal marker (catalase) confirmed the absence of peroxisomes in PEX3 KO or PEX19 KO HEK293T cells. Scale bar: 10 mm. (c) Schematic view of the constructs used in glycosylation assays. (d) Intracellular localization of the constructs used in glycosylation assay (e) analyzed by immunofluorescence. HEK293T cells were transiently transfected with indicated constructs for 24 h. Calnexin was used to label ER, catalase was used to label peroxisomes (PO). Scale bar: 10 mm. (e) TA substrates with or without OPG tag were transfected to WT, PEX3 KO or PEX19 KO HEK293T cells. Glycosylation bands are indicated by asterisk. Glycosylation was confirmed by loss of the band after addition of glycosidase, PNGase-F

To test whether peroxisomal TA proteins insert into the ER on their way to peroxisomes, we used a previously described model protein YgiM(TA), a bacterial derived sequence, shown to specifically target to peroxisomes in mammalian cells ^20^. As a reporter for protein entry into the ER we used opsin glycosylation tag (OPG) since it contains a glycosylation site which accepts glycans if exposed to the ER lumen ^21^. OPG was fused to the C-terminus of YgiM(TA), a peroxisomal TA construct, or SEC61β, a known ER-targeted TA construct (Fig 1c). Confocal microscopy coupled with specific immunostaining demonstrated that the OPG tag did not influence localization of either construct (Fig. 1d).

WT, PEX3 KO or PEX19 KO HEK293T cells were transiently transfected with peroxisomal TA, YgiM(TA)-OPG, or ER-targeted TA, SEC61β-OPG constructs. SEC61β-OPG expressed in control cells or cells lacking peroxisomes showed clear glycosylation as detected by a western blot band shift when treated with PNGase-F to remove oligosaccharides from glycoproteins (Fig. 1e). In contrast, YgiM(TA)-OPG glycosylation was non-detectable in control cells or in cells lacking peroxisomes. Together, these results suggest that peroxisomal TA proteins do not enter ER before their insertion into the peroxisome. Also, peroxisomal TA proteins do not appear to target to the ER even in the absence of mature peroxisomes (Fig. 1e).

### Peroxisomal TA proteins are targeted to mitochondria in the absence of peroxisomes

Since peroxisomal TA proteins do not appear to enter ER in control cells or cells lacking peroxisomes, we sought to identify the localization of peroxisomal TA proteins in cells devoid of peroxisomes. For this purpose, we generated cells stably expressing peroxisomal TA, YgiM(TA), in WT, PEX3 KO or PEX19 KO HEK293T cells. We analyzed the colocalization of YgiM(TA) with peroxisomes, ER or mitochondria by immunofluorescence microscopy and quantitative colocalization analysis. In WT cells YgiM(TA) localized strongly to peroxisomes (Fig. 2 a-c). In PEX3 KO or PEX19 KO cells YgiM(TA) showed colocalization with both ER and mitochondria, with most notable colocalization with mitochondria (Fig. 2c).

**Fig. 2.**
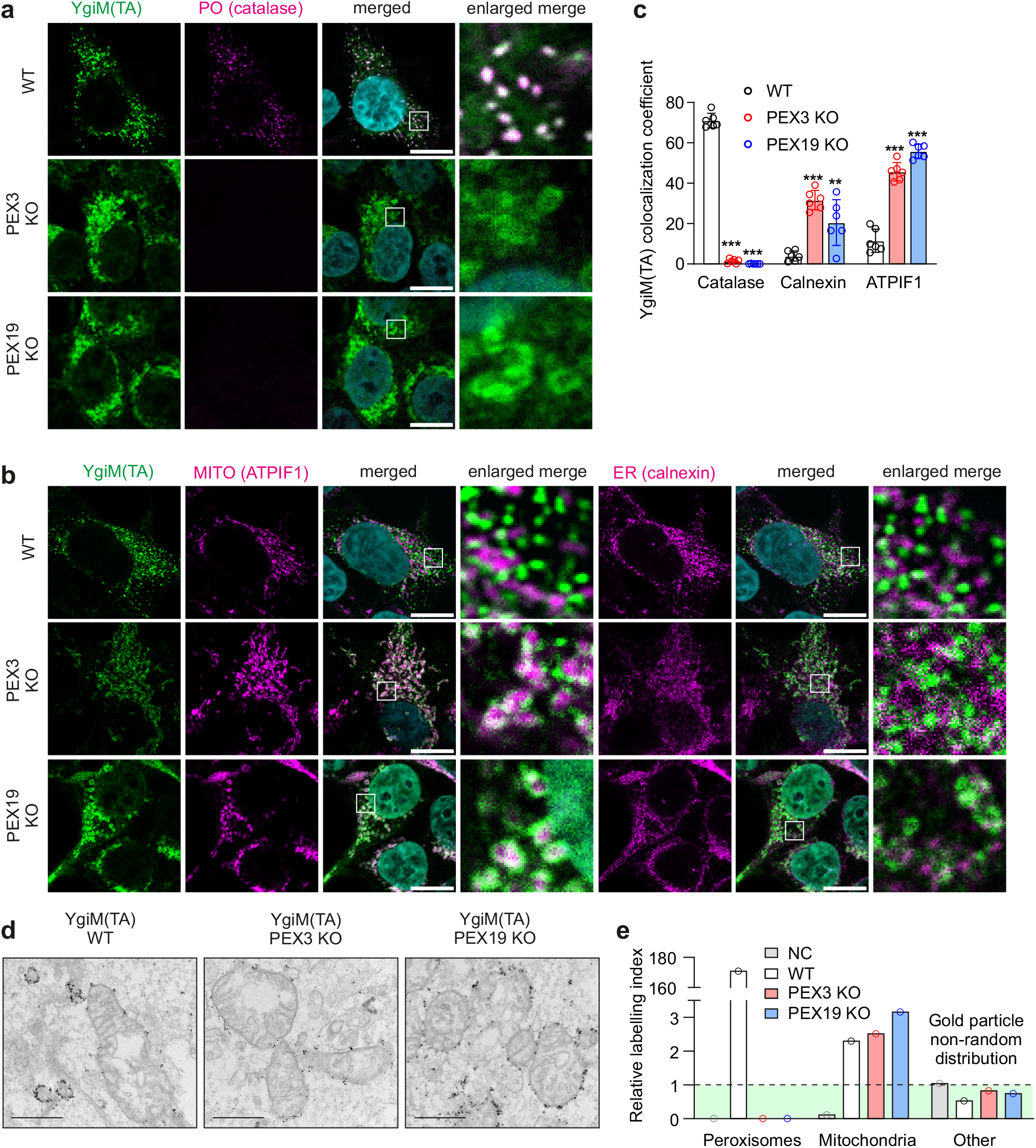
Exogenous peroxisomal TA YgiM(TA) is targeted to mitochondria in mammalian cells lacking peroxisomes. Localization of EGFP-YgiM(TA) stably expressed in WT, PEX3 KO or PEX19 KO HEK293T cells was analyzed by immunofluorescence confocal microscopy. (a) Catalase was used to label peroxisomes (PO). (b) ATPIF1 was used to label mitochondria (MITO) and calnexin was used to label ER. Scale bar, 10 mm. (c) Quantification analysis of confocal microscopy images by Mander’s overlap coefficient. The data are presented as mean ± SD. **P < 0.01, ***P < 0.001 as compared to WT cells (unpaired t-tests). (d) WT, PEX3 KO or PEX19 KO HEK293T cells stably expressing EGFP-YgiM(TA) were immunolabeled with anti-GFP antibody for transmission electron microscopy. (e) Stereological analysis of electron microscopy images was used to identify relative labeling index of YgiM(TA) in peroxisomes, mitochondria or other organelles. NC, negative control, cells not expressing EGFP-YgiM(TA) construct used for quantitation of background signal from immunogold labeling. Area above the green surface depicts non-randomly distributed gold particles. Analyzed area of 300 μm2, 2 images per cell, 15 cells total.

To visualize YgiM(TA) localization at subcellular level, we used immuno-electron microscopy. Immunogold electron microscopy analysis showed that while in WT cells YgiM(TA) localized mainly to peroxisomes in the absence of peroxisomes it was primarily targeting mitochondria (Fig. 2d). We analyzed electron microscopy images using stereological analysis to identify mitochondrial relative labeling index ^22, 23^. Consistent with the immunofluorescence experiments (Fig. 2 a-c), we observed robust localization of YgiM(TA) to mitochondria in PEX3 KO or PEX19 KO cells (Fig. 2e).

YgiM(TA) is an exogeneous peroxisomal TA protein that was overexpressed in HEK293T cells in our model. Since overexpressed proteins have the propensity to mistarget we analyzed localization of an endogenous peroxisomal TA protein, ACBD5, in cells lacking peroxisomes. By immunofluorescent microscopy and quantitative colocalization analysis we showed that ACBD5 is strictly localized to peroxisomes in WT HEK293T cells (Fig. 3 a, d). In PEX19 KO cells ACBD5 was detected in mitochondria, but not in the ER (Fig. 3 b-d). Notably, ACBD5 was barely detectable by immunofluorescent analysis or by immunoblotting in PEX3 KO cell (Fig. 3 a-c, e), suggesting its degradation. Thus, endogenous peroxisomal TA proteins localize to mitochondria in the absence of peroxisomes, suggesting that mitochondria, but not ER may be involved in peroxisomal targeting of TA proteins.

**Fig. 3.**
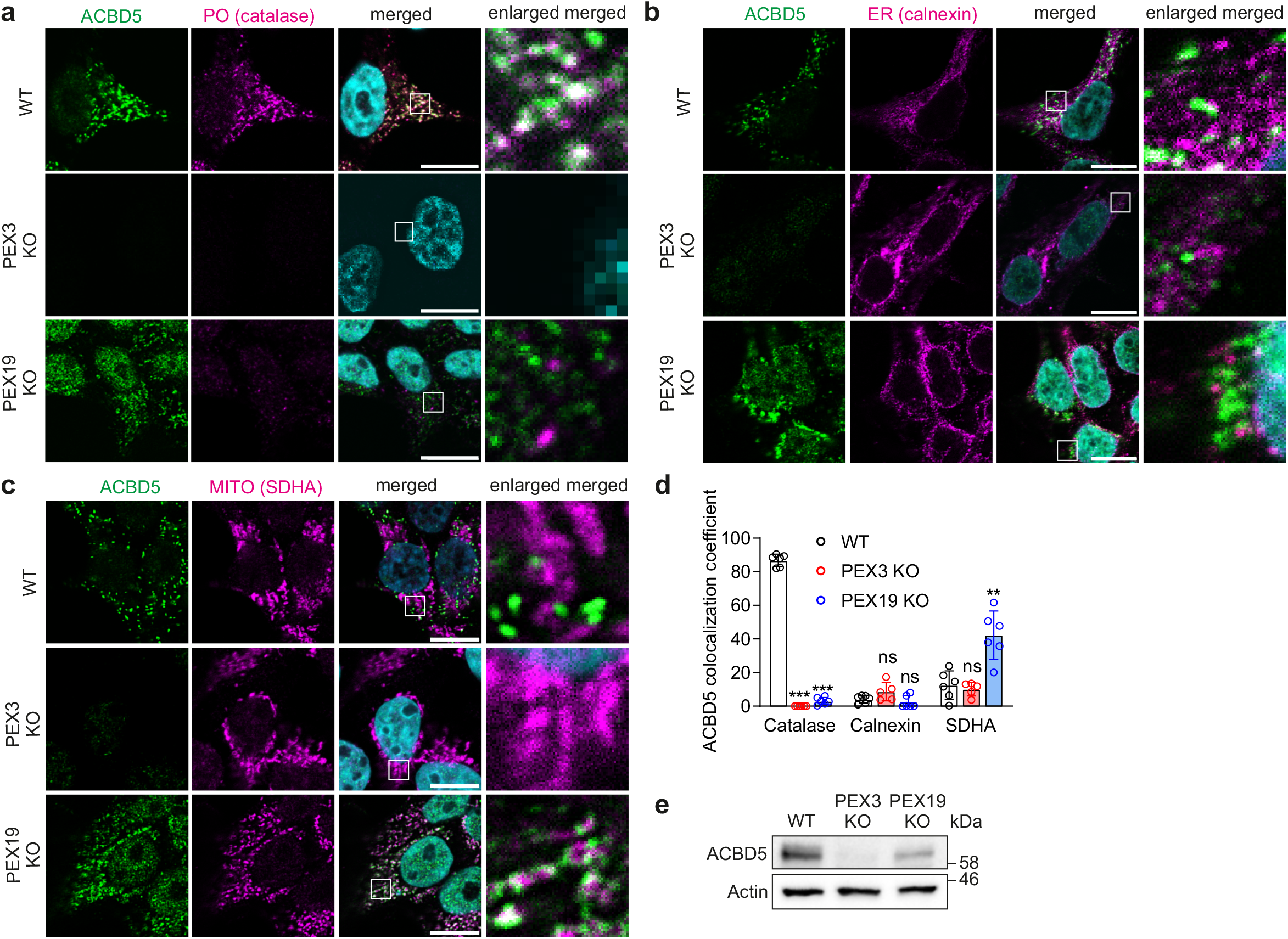
Endogenous peroxisomal TA protein ACBD5 localizes to mitochondria in the absence of peroxisomes in mammalian cells. (a-c) Localization of ACBD5 in WT, PEX3 KO or PEX19 KO HEK293T cells was analyzed by immunofluorescence confocal microscopy. (a) Catalase was used to label peroxisomes (PO). (b) Calnexin was used to label ER. (c) SDHA was used to label mitochondria (MITO). Scale bar, 10 mm. (d) Quantification analysis of confocal microscopy images by Mander’s overlap coefficient. The data are presented as mean ± SD. **P < 0.01, ***P < 0.001, ns, not significant as compared to WT cells (unpaired t-tests). (e) Immunoblotting analysis of ACBD5 in WT, PEX3 KO or PEX19 KO HEK293T cells. Actin was used as a loading control.

### Peroxisomal TA proteins insert to mitochondria *en route* to the peroxisome in mammalian cells

Having established that in the absence of peroxisomes, peroxisomal TA proteins localize to mitochondria, we sought to investigate the route by which they reach peroxisomes under normal conditions. To achieve this, we used a proximity biotinylation approach. We used wild type BirA/AviTag labeling approach ^24^ to reveal interactions between peroxisomal TA (AviTagged) and specific organelles containing a localized BirA protein. This approach is based on the ability of a biotin ligase (BirA) to biotinylate AviTag in a highly specific manner only if both tagged proteins are present in the same cellular compartment ^25, 26^. We introduced a doxycycline-inducible BirA construct targeting ER (ER-BirA), mitochondrial intermembrane space (IMS-BirA) or peroxisomes (PO-BirA) in the Flp-In T-Rex™ HEK293 cells using corresponding targeting sequences. We then fused the AviTag sequence to the C-terminus of EGFP-tagged ER or peroxisomal TA protein (Fig. 4 a,b). These AviTag constructs were subsequently transiently transfected to the Flp-In T-Rex™ HEK293 cells stably expressing different organelle-targeted BirA constructs. To avoid overexpression of the AviTag constructs, we used the weak promoter, UBC ^27^. We then confirmed the expected organelle localization of BirA and AviTag constructs using immunofluorescence microscopy (Fig. 4c). To ensure the specificity of the assay we kept the expression of BirA at a very low level by using inherent leakiness in our doxycycline system without adding additional doxycycline to the culture media.

**Fig. 4.**
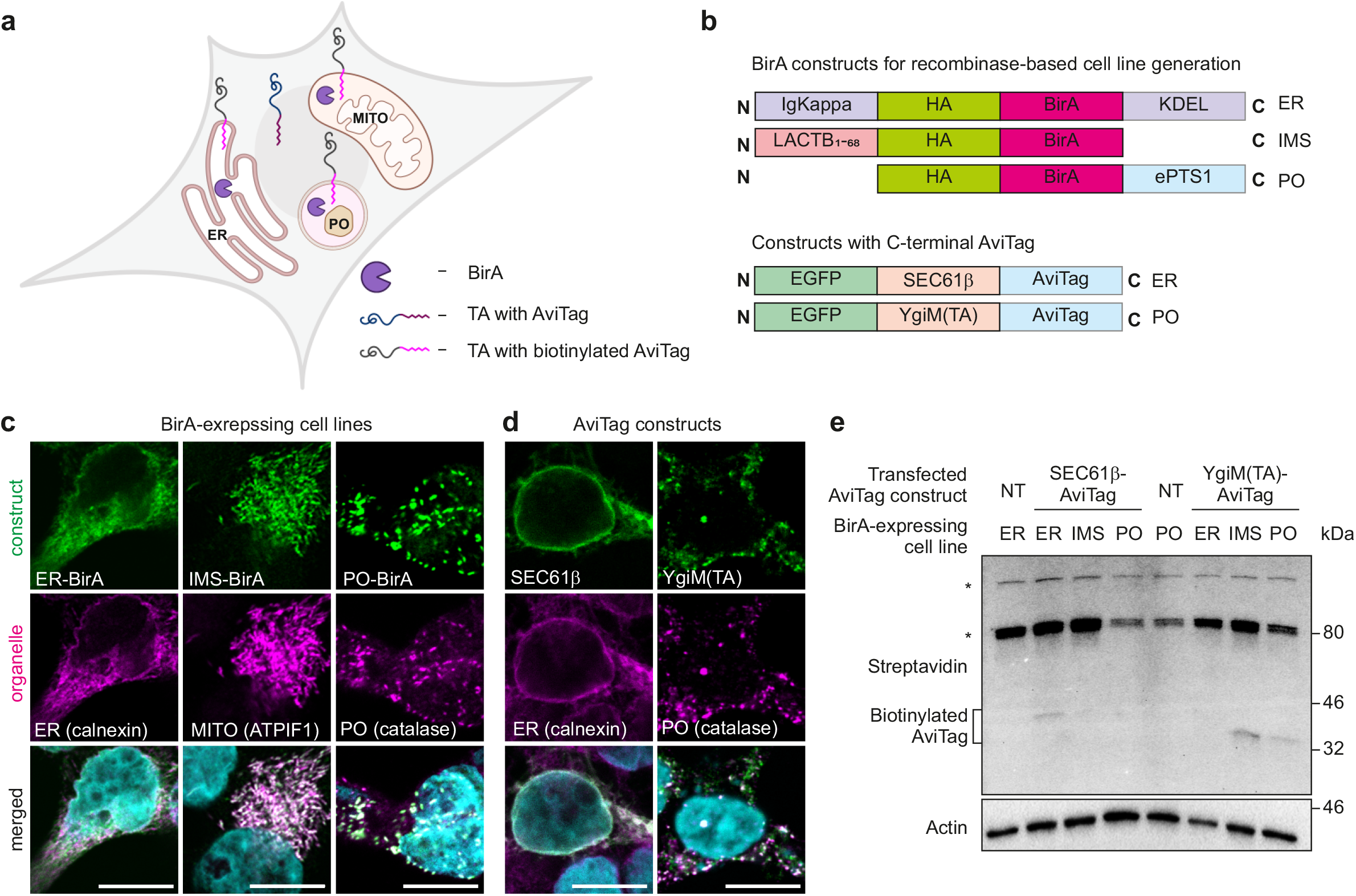
Peroxisomal TA pass through mitochondria on the way to peroxisome in mammalian cells. (a) Schematic representation of the biotinylation experiment. TA constructs fused to AviTag are transfected to the cells expressing BirA in endoplasmic reticulum (ER), peroxisomes (PO) or mitochondria (MITO). Once TA inserts to the specific organelle labeled with BirA, biotin acceptor sequence of the TA, AviTag, is biotinylated by BirA. (b) Schematic view of the constructs used to generate BirA-expressing cell lines and TA constructs fused to AviTag. (c) Intracellular localization of doxycycline inducible BirA constructs targeted to the specific organelle in Flp-In™ T-REx™-293 cells was analyzed by immunofluorescence microscopy. ER, endoplasmic reticulum, IMS, mitochondrial intramembrane space, PO, peroxisomal matrix. Scale bar, 10 mm. (d) Intracellular localization of AviTag-containing TA constructs transiently transfected to Flp-In™ T-REx™-293 cells was analyzed by immunofluorescence confocal microscopy. Scale bar, 10 mm. (e) Representative streptavidin immunoblot of BirA-expressing cells transfected with ER targeted TA, SEC61β-AviTag or peroxisomal targeted TA, YgiM(TA)-AviTag. Bands of ∼33 kDa and 39 kDa indicate biotinylated AviTag constructs. Endogenous biotinylated mitochondrial proteins are shown by asterisks. Actin was used as a loading control. NT, non-transfected control. Figure (a) was created with BioRender.com

Flp-In T-Rex™ HEK293 cells expressing ER-BirA, IMS-BirA or PO-BirA were transiently transfected with ER-targeted TA construct, SEC61β-AviTag, or with peroxisome-targeted TA construct, YgiM(TA)-AviTag. Using streptavidin immunoblotting we showed that as expected SEC61β-AviTag was biotinylated in ER-BirA cells but not in IMS-BirA or PO-BirA expressing cells (Fig. 4e), validating the assay as a reliable tool to analyze the route of targeting of TA proteins that are ultimately delivered to different target organelles. In contrast, YgiM(TA)-AviTag biotinylation was exclusively detected in cells expressing PO-BirA or IMS-BirA (Fig. 4e). Therefore, we conclude that peroxisomal TA proteins pass through mitochondria, but not through ER, in cells containing intact mature peroxisomes.

Collectively, our results showed that peroxisomal TA proteins target mitochondria in mammalian cells both in the presence and in absence of peroxisomes, suggesting that mitochondria are involved in the targeting of peroxisomal TA proteins.

## DISCUSSION

Research on targeting pathways of peroxisomal TA in mammalian cells provided data both in favor ^7^ and against a route involving the ER ^28, 29^. It has been demonstrated that in yeast ^30, 31^ and plants ^32^ peroxisomal TA proteins insert to ER prior to reaching peroxisome. Further, in yeast cells devoid of peroxisomes, peroxisomal TA proteins were visualized in what were described as pre-peroxisomal vesicles derived from ER ^20, 33–35^. Using glycosylation assays in human cells, we tested the possibility of peroxisomal TA transiting through the ER in both WT and cells lacking peroxisomes. To generate cells lacking peroxisomes we knocked out essential membrane or cytosolic peroxisomal biogenesis factors, PEX3 or PEX19, respectively. The absence of N-linked glycosylation provided evidence against ER-dependent targeting of peroxisomal TA proteins in mammalian cells. However, we cannot fully exclude the possibility that peroxisomal TA proteins might transiently visit the ER lumen, without becoming glycosylated. Therefore, we analyzed the localization of peroxisomal TA proteins in mammalian cells by immunofluorescence and immuno-electron microscopy imaging. We observed that in the absence of peroxisomes in PEX19 KO cells, endogenous peroxisomal TA protein ACBD5 localizes mainly to mitochondria. Mitochondrial localization of peroxisomal membrane proteins in the absence of peroxisomes has been reported previously ^36, 37^, although, this finding was considered as a mistargeting artifact. However, we showed that in PEX3 KO cells, which lack peroxisomes as well, ACBD5 does not target mitochondria. These data suggest that mitochondrial localization of peroxisomal membrane proteins may not represent simple passive mistargeting, but instead an active process in which PEX3 may play an important role. Interestingly, PEX3 and PEX14 were found to be targeted to mitochondria prior to their assembly into peroxisomes (Sugiura et al) ^8^, suggesting the involvement of these factors in the insertion of peroxisomal TA proteins to mitochondria. We observed almost complete depletion of ACBD5 in PEX3 KO cells, suggesting that at least some untargeted peroxisomal TA proteins become directed for degradation. Notably, PEX19 KO causes reduction of PEX3 protein level, suggesting that being a peroxisomal membrane protein, PEX3 undergoes degradation in the absence of peroxisomes. However, the residual level of PEX3 in PEX19 KO cells was sufficient to protect ACBD5 from complete degradation in PEX19 KO cells.

Using organelle-specific proximity biotinylation assays we showed that peroxisomal TA proteins target both peroxisomes and mitochondria in cells containing intact peroxisomes. This indicates that mitochondria-related targeting is a process that normally occurs during peroxisomal TA trafficking under normal conditions. Overall, our data suggest that TA targeting to mitochondria, previously described as mislocalization, may be part of the endogenous targeting pathway to peroxisomes. These observations are consistent with the findings by Sugiura et al. ^8^ showing that newly assembled peroxisomes arise from both ER and the mitochondria.

The results obtained in our study are still to be further confirmed if they are to be generalized across all known peroxisomal TA, and the routes that specific TA proteins follow to reach peroxisome remain to be examined. We expect future studies to illuminate the precise delivery route of peroxisomal TA proteins and detailed kinetics of their mitochondrial passage. Optogenetic approaches with organelle-targeted photoactivatable fusion proteins have revealed details of protein targeting to other cellular organelles ^38^ so will also likely be useful for revealing details of the mitochondria-peroxisome delivery pathway of peroxisomal TA proteins.

## MATERIALS AND METHODS

### Cell culture

Human embryonic kidney cells (HEK293T) or Flp-In T-REx™-293 cells (ThermoFisher, R780-07) were cultured (37°C, 5% CO2) in DMEM (Pan-Biotech, P04-03600), supplemented with 10% fetal bovine serum (GIBCO, 10270106), l-glutamine (GIBCO, 25030081) and penicillin/streptomycin (GIBCO,15140122). TransIT-X2 (MirusBio, MIR 6000) was used for cell transfection according to manufacturer’s instruction.

### Generation of knockout cell lines

CRISPR/Cas9 was used to generate knockout HEK293T cells. Cells were co-transfected with two gRNAs and pSpCas9n(BB)-2A-Puro (PX462) V2.0 (Addgene, 62987). After 24 h the cells were selected with puromycin and expanded. T7 assay was used to confirm genome editing. Single cell clones were obtained and validated using immunoblotting analysis.

### Generation of stably expressing cell lines

HEK293T cells stably expressing EGFP-YgiM(TA) were generated by using standard lentiviral cell line generation protocol. Briefly, WT, PEX3 KO or PEX19 KO cells were co-transfected with TransIT-2020 (MirusBio, MIR 5400), pMD2.G (Addgene, 12259), psPAX2 (Addgene,12260) and pLL3.7-EGFP-YgiM(TA) following manufacturer’s instructions. Single cell clones were selected for medium expression by FACS.

Doxycycline-inducible BirA constructs was introduced into HEK293T by co-transfecting Flp-In™ T-REx™-293 with pOG44 (ThermoFischer, V600520) and corresponding BirA construct using TransIT-2020 (MirusBio, MIR5400) according to manufacturer’s instructions.

### Plasmid construction

OPG-taged plasmids were constructed by homologous recombination in yeast as previously described ^39^ . Briefly, to generate pEGFP-SEC61β-OPG, mCherry-SEC61β was PCR amplified from mCh-Sec61 beta (a gift from Gia Voeltz (Addgene plasmid, 49155; http://n2t.net/addgene:49155; RRID:Addgene 49155)) with primers ATGGTGAGCAAGGGCGAGGA and TTACCCTGTCTTATTGCTAAATGGAACGTAAAAGTTAGGACCCGAACGAGTGTACTTGCCCCAAATGTG. SEC61β-OPG was generated by PCR amplification using primers ATCACTCTCGGCATGGACGAGCTGTACAAGAGATCTATGCCTGGTCCGACC and GGTATGGCTGATTATGATCAGTTATCTAGATTACCCTGTCTTATTGCTAAATGGA AC and the obtained product was co-transformed with pEGFP-C1-2µ-URA3 cut with BamHI (NEB, R3136) into yeast strain BY4743 (EUROSCARF Y20000) cut. The obtained yeast plasmid DNA transformed into competent cells (NEB, C2987H).

To generate pEGFP-YgiM(TA)-OPG, YgiM(TA)-OPG sequence was produced by PCR from pRS315-mCherry-YgiM(TA) with primers ATCACTCTCGGCATGGACGAGCTGTACAAGATGGTAGAGGATAAGATCCAGAAGGAAACA and TATGGCTGATTATGATCAGTTATCTAGATTACCCTGTCTTATTGCTAAATGGAACGTAAAAGTTAGGACCGTTCATCCAGCGATCTTTG. The resulted PCR product co-transformed with pEGFP-C1-2µ-URA3 cut with BamHI (NEB, R3136) into yeast strain BY4743 (EUROSCARF Y20000) cut. The obtained yeast plasmid DNA was transformed into competent cells (NEB, C2987H). Yeast strains, plasmids, and culture conditions are as described in ^39^.

To generate pLL3.7-EGFP-YgiM(TA), pLL3.7 (Addgene, 11795) was cut with EcoRI (NEB, R3101) and NheI (NEB, R3131) and the 6907 bp band was gel purified (Macherey-Nagel, 740609.50). EGFP-YgiM(TA) was PCR amplified from pEGFP-YgiM(TA) with primers ATATTGCTAGCGCTACCGGTCGCCACCATG and CCTACTGAATTCTTAGTTCATCCAGCGATCTTTGC. The obtained product was then cut with NheI (NEB, R3131) and EcoRI (NEB, R3101) and PCR purified (Macherey-Nagel, 740609.50). The two purified fragments were then ligated using T4 DNA ligase (NEB, M0202) following manufacturer’s instructions and transformed into competent cells (NEB, C2987H).

To generate BirA-constructs, pDisplay-BirA-ER (a gift from Alice Ting (Addgene plasmid # 20856; http://n2t.net/addgene:20856 ; RRID:Addgene_20856)) was used as a PCR template for ER-BirA or PO-BirA. The HA-BirA region was PCR amplified with the following primers: for ER-BirA: GCGAAGCTTTGGGGATATCCACCATGGAGACAGAC/GCGGGATCCGTTCGTCGACTCACAGCTCGTCCTT, for PO-BirA: TTTTAAGCTTGCCACCATGTATCCATATGATGTTCC/TTTTGGATCCTCATAGCTTACTTCTTCTGCCACGCCCCAGTTTTTCTGCACTAC. IMS-BirA (LACTB_1-68_-HA-BirA) construct was purchased from Twist Bioscience (San Francisco, California, United States). The IMS sequence was as in ^40^. The IMS-BirA construct and PCR products from ER-and PO-BirA products were cut with HindIII (NEB, R3104) and BamHI (NEB, R3136) and cloned into pcDNA™5/FRT/TO (TermoFischer, V652020) that was cut with the same restriction enzymes.

To generate AviTag constructs, pLJC5-3XHA-EGFP-PEX26 (Addgene, 139054) was cut with AgeI (NEB, R3552) and EcoRI(NEB, R3101). The large DNA fragment was gel purified (Macherey-Nagel, 740609.50) and used as a target backbone for cloning the AviTag constructs. AgeI-EGFP-SEC61β sequence and linker AviTag-EcoRI were purchased from Integrated DNA Technologies (IDT, Coralville, Iowa, USA), PCR amplified, annealed, ligated with the cut backbone and transformed into competent cells (NEB, C2987H). EGFP-YgiM(TA) was PCR amplified from pEGFP-YgiM(TA)-noFis1(119-128) with appropriate primers, annealed with linker AviTag sequence as above and cloned to competent cells. GSGSTSGSGK was used as a linker sequence ^24^.

### Glycosylation assay in cells

Cells were transiently transfected with pEGFP-SEC61β-OPG or pEGFP-YgiM(TA)-OPG using TransIT2020 transfection reagent (MirusBio, MIR 5400) according to manufacturer’s instructions. After 24 h cell lysates were treated with PNGase-F (NEB, P0704L) or water by following the manufacturer’s protocol. The samples were resolved by SDS-PAGE and immunoblotted.

### Immunofluorescence analysis

HEK293T or Flp-In™ T-REx™-293 were fixed with 4% PFA for 10 min on 13 mm coverslips and then washed with PBS three time. Next the cells were permeabilized with 1% TritonX-100 (CAS 9002-93-1) for 15 min and then DAPI (1:1000 in PBS) was added for 10 min. The coverslips were washed three rimes with PBST (PBS containing 0.1% Tween20 (Fisher Scientific, 10485733)) and blocked with 1% BSA (Fisher Scientific, 11413164) in PBST for 2 h. Cells were then incubated with corresponding primary antibody diluted in blocking buffer at 4°C overnight. After washing for 15 min three times, the corresponding secondary antibodies were added for 1 h at room temperature. After washing for 5 min three times the samples were mounted with antifade mounting medium (Vector Laboratories, H-1700). The samples were imaged on Leica TCS SP8 STED 3X CW 3D microscope using HC PL APO 93x/1.30 motCORR STED WHITE (glycerol, wd 0.3mm) objective. Primary antibodies used for immunofluorescence analysis: rabbit anti-ACBD5, 1:300 (Atlas Antibodies, HPA012145), chicken anti-GFP, 1:10000 (Aves, GFP-1020), rabbit anti-catalase, 1:400 (CST, 12980S), rabbit anti-calnexin, 1:1000 (Abcam, 22595), rabbit anti-ATPIF1, 1:250 (CST, 8528), mouse anti-SDHA, 1:100 (Santa Cruz, 390381), mouse anti-calnexin, 1:500 (Abcam, 112995), mouse anti-HA, 1:400 (Santa Cruz, 2367), mouse anti-catalase 1,6 µg / mL (Biotechne, MAB3398). Secondary antibodies: Alexa Fluor 647 (Invitrogen, A-21235 or CST, 4414S), Alexa Fluor 555 (CST, 4413S or CST, 4409S), Alexa Fluor 488 (Invitrogen, A-11001 or CST, 4412S).

### Colocalization analysis

Colocalization analysis was performed with the Imaris Software (Imaris File Converter 9.5.1) using the Coloc tool of the Surpass system. Images were manually adjusted to separate background from the real signal. Mander’s overlap coefficient was used to determine the percentage overlap. For each sample 5-6 fields of view where each field of view captured approximately 50 cells were analyzed.

### Western blotting

Cells were washed with PBS and lysed in RIPA buffer (CST, 9806). Protein concentration was determined by BCA assay kit (Thermo Scientific, 23227). Protein lysates were supplemented with Laemmli buffer, heated at 95°C for 5 min and separated by SDS-PAGE. Then the proteins were transferred to a PVDF membrane using Trans-Blot Turbo Mini 0.2 µm PVDF Transfer Pack (BioRad, 1704156). The membranes were blocked in 5% non-fat milk in TBST for 1 h at room temperature, incubated with appropriate primary antibodies diluted in 1% BSA in TBST for 1 h at RT or overnight at 4°C. Next the membranes were washed three times in TBST followed by 1 h incubation at room temperature with appropriate secondary antibodies diluted in 1% BSA in TBST, washed trice in TBST and imaged for either chemiluminescence or IR signal. Primary antibodies: rabbit anti-GFP 1:2500 (Cell Signaling Technology, 2956), rabbit anti-PEX19 1:1000 (Abcam, 137072), rabbit anti-ACBD5 1:500 (Atlas Antibodies, HPA012145), mouse anti-HA 1:1000 (Cell Signaling Technology, 2367), mouse anti-β-actin, 1:1000 (Cell Signaling Technology, 3700), IRDye 800CW conjugated streptavidin 1:3000 (LI-COR, 926-32230). Secondary antibodies: anti-mouse IgG HRP-linked antibody 1:5000 (Cell Signaling Technology, 7076). The chemiluminescence was developed using ECL substrate for enhanced chemiluminescence (Thermo Scientific, 32106) and the signal was captured by Chemidoc imaging system (Bio-Rad). IR signal was detected using iBright Imaging Systems (Invitrogen).

### Immuno-electron microscopy

WT, PEX3 KO or PEX19 KO HEK293T cells were fixed with paraformaldehyde-lysine-periodate ^41^, permeabilized with 0.01% saponin (Sigma-Aldrich, S7900) and stained for 1.5 h with anti-GFP antibody (1:500, Sigma, 290), followed by 1 h incubation with 1.4 nm nanogold-conjugated secondary antibody (1:60, RRID: AB_2631182, NY, Nanoprobes, Stony Brook, 2004). Silver enhancement of nanogold particles was performed according to manufacturer’s instructions with HQ Silver Kit (Nanoprobes, Stony Brook, NY), then gold toned with 2% Na-Acetate, 0.05% HAuCl4, and 0.3% Na2S2O3⋅5H2O. Post-fixation was done for 1h on ice with 1% reduced osmium tetroxide in sodium cacodylate buffer pH 7.4, dehydrated by serial washing in ethanol and acetone and finally embedded in Epon (TAAB 812, Aldermaston, UK) for 2 h before 14 h polymerization at 60°C. Sections were prepared with Leica UCT6 microtome, then post-stained with lead citrate and uranyl acetate. Samples were imaged with Jeol JEM-1400 (Jeol Ltd., Tokyo, Japan) operated at 80 kV, with Orius CCD-camera Gatan SC 1000B, bottom mounted (Gatan Inc., Pleasanton, CA). Quantification of the relative labeling index was performed as previously described ^42^, software used was Microscopy Image Browser ^43^ to segment and calculate.

### Biotinylation assay

Flp-In™ T-REx™-293 cells were grown in 6 well-plates and transiently transfected with 250 ng of EGFP-SEC61β-AviTag or 1000 ng of EGFP-YgiM(TA)-AviTag for 24 h and processed for immunoblotting. Membranes were incubated with IRDye 800CW conjugated streptavidin.

### Statistical analysis

All data are presented as mean ± standard deviation (SD). For the statistical analysis with two samples, unpaired two-tailed t-test was performed using Graph Pad Prism 9 Software. All the graphs were prepared with Graph Pad Prism 9 Software.

## ACKNOWLEDGEMENTS

We gratefully acknowledge grant support provided to Cory Dunn from ERC Starting Grant (grant nr. 637649), Academy of Finland, Finland (grant nr. 331556), Jane ja Aatos Erkon Säätiö, Finland (grant nr. 200057) and Sigrid Jusèlius Foundation, Finland.

We thank Pekka Katajisto for valuable discussion and suggestions, members of Ville Paavilainen laboratory for discussions and insights throughout this project, Leonardo Souza-Almeida and Maria Vartiainen for continuous support.

## AUTHOR CONTRIBUTIONS

T.S.: experimental tools and resources generation, methods, study design, investigation, analysis, imaging, manuscript writing, figure preparation. G.L.B: glycosylation in cells assay design, imaging and preparation. F.K.: initial optimization of biotinylation experiment conditions. H.V.: immuno-electron microscopy sample preparation, imaging, and analysis.

A.P.: mentoring and guidance throughout the project and participation in planning the design of BirA constructs. E.J: discussion on the analysis of the IEM data. V.P: supervision of the project. S.K.: supervision and guidance of the project, manuscript writing. All authors have read and agreed to the published version of the manuscript.

